# The effects of locomotion on sensory-evoked haemodynamic responses in the cortex of awake mice

**DOI:** 10.1101/2021.12.16.472794

**Authors:** Beth Eyre, Kira Shaw, Paul Sharp, Luke Boorman, Llywelyn Lee, Osman Shabir, Jason Berwick, Clare Howarth

**Author notes:** These authors contributed equally (Beth Eyre, Kira Shaw). These authors contributed equally (Jason Berwick, Clare Howarth).

## Abstract

Investigating neurovascular coupling in awake rodents is becoming ever more popular due, in part, to our increasing knowledge of the profound impacts that anaesthesia can have upon brain physiology. Although awake imaging brings with it many advantages, we still do not fully understand how voluntary locomotion during imaging affects sensory-evoked haemodynamic responses. In this study we investigated how evoked haemodynamic responses can be affected by the amount and timing of locomotion. Using an awake imaging set up, we used 2D-Optical Imaging Spectroscopy (2D-OIS) to measure changes in cerebral haemodynamics within the sensory cortex of the brain during either 2s whisker stimulation or spontaneous (no whisker stimulation) experiments, whilst animals could walk on a spherical treadmill. We show that locomotion alters haemodynamic responses. The amount and timing of locomotion relative to whisker stimulation is important, and can significantly impact sensory-evoked haemodynamic responses. If locomotion occurred before or during whisker stimulation, the amplitude of the stimulus-evoked haemodynamic response was significantly altered. Therefore, monitoring of locomotion during awake imaging is necessary to ensure that conclusions based on comparisons of evoked haemodynamic responses (e.g., between control and disease groups) are not confounded by the effects of locomotion.

## Introduction

When neurons fire, there follows a localised increase in blood flow to that same brain region. This relationship between neuronal firing and an increase in blood flow is known as neurovascular coupling (NVC), and underpins the principles of blood oxygen level dependent functional magnetic resonance imaging (BOLD fMRI). NVC ensures that the brain receives prompt increases in cerebral blood flow (CBF) to activated regions of the brain, allowing for the rapid delivery of essential nutrients such as O_2_ and glucose and the removal of waste products such as CO_2_ and lactate^1^. This mechanism is important for healthy brain function; accumulating evidence suggests that NVC is impaired in several neurological disorders, including Alzheimer’s disease^2-5^. Therefore, understanding how NVC may be altered by disease is integral to furthering our understanding of the onset and progression of such diseases.

Over the last fifty years, the neurovascular field has been revolutionised by the advent of new scientific methods. The development of techniques such as wide-field optical imaging^6,7^ and two-photon microscopy^8^, alongside advances in the development of genetically-encoded calcium indicators such as GCAMP^9^, have been vital in furthering our knowledge of the inner workings of the brain. Just considering the last twenty years, we have discovered the potential role of pericytes^10-12^, astrocytes^13-15^ and even caveolae^16^ in NVC. Many studies investigating NVC, including several of those previously mentioned were conducted under anaesthesia. The use of anaesthesia has allowed the field to gain an in-depth insight into neural activity and the subsequent haemodynamic response in a controlled environment. However, anaesthesia is not without its pitfalls^17^.

Not only does anaesthesia dampen neural activity but it can also reduce many aspects of the haemodynamic response - including blood oxygenation, CBF and cerebral blood volume (CBV)^18,19^ in addition to delaying the time-course of the haemodynamic response^18^.

To mitigate these effects, several groups, including our own, have developed anaesthetic regimes that produce stable haemodynamic responses, of similar timing and magnitude to those in the awake preparation^20-22^ - these regimes can produce stable responses without the confounds of behaviour. Despite this, many groups have begun to move away from the use of anaesthesia. A growing number of studies have used awake, moving animals to investigate NVC and the roles of cells within the neurovascular unit (NVU) ^16,23,24^.

Studies using electrophysiology, two-photon microscopy and intrinsic optical signal (IOS) imaging have shown that locomotion can generate robust increases in neural activity, vessel diameter, and CBV, respectively^23,25,26^. While these studies focused on how locomotion itself impacts brain haemodynamics (within surface vessels of the brain), Tran et al. (2018) explored how locomotion may affect sensory-evoked haemodynamic responses. No significant differences in peak amplitude dilation of arterioles were reported when mice were continuously running during whisker stimulation, went from quiet to running in response to whisker stimulation, or when they remained quiet prior to and after whisker stimulation^24^ – suggesting that locomotion did not have an impact on sensory-evoked haemodynamic responses. However, the study focused on dilation changes in penetrating arterioles, and locomotion may affect sensory-evoked haemodynamic responses within the surface vessels of the brain to a different extent, as it has been shown that the narrowing of the Virchow-Robin space may restrict dilations within penetrating arterioles^27^.

It is critical to characterise how sensory-evoked haemodynamic responses may be impacted by locomotion for a number of reasons. If sensory-evoked haemodynamic responses are affected by locomotion and locomotion is not monitored, when comparing responses (for example when comparing a disease group to a wild-type (WT) group), differences in neurovascular function could be erroneously assumed to be a consequence of disease, rather than a consequence of differential locomotion.

Therefore, to improve our understanding of how locomotion may impact sensory-evoked haemodynamic responses we used two-dimensional optical imaging spectroscopy (2D-OIS) to investigate changes in cortical blood oxygenation in C57BL/6J mice. We hypothesised that locomotion would increase the amplitude of evoked haemodynamic responses within the cerebral cortex of the brain, as previous research suggests that locomotion leads to greater dilations within surface vessels as compared to penetrating vessels^27^. Additionally, we hypothesised that the time at which locomotion occurred (in relation to the 2s whisker stimulation) would also impact evoked haemodynamic responses, with locomotion occurring closer to the stimulation onset being expected to increase the amplitude of the evoked hemodynamic response. The data set included 21 separate imaging sessions taken from 4 animals, with each session comprised of continuous recordings taken during (a) a 2s whisker stimulation experiment (59 × 25-second trials, with whisker stimuli presented between 5-7s after trial onset) and (b) a spontaneous experiment, with no whisker stimulation (also separated into 59 × 25-second trials). For the 2s whisker stimulation data set recorded in each of the individual 21 sessions, the following analysis was conducted. The 59 trials were ranked by the amount of voluntary locomotion occurring during the entire trial period (25s), and during different 5s time windows within the trial (occurring before, during or after stimulation). Average evoked haemodynamic time series were created from the top (n=6 trials per session) and bottom (n=6 trials per session) 10% of ranked trials, corresponding to those with the most and least locomotion respectively, this was completed for each session. For visualisation and analysis purposes, average evoked time series were then generated by averaging across all 21 sessions (giving mean +/-SEM between sessions). From this analysis, we were able to reveal how different amounts of locomotion impact the amplitude of the sensory-evoked haemodynamic response, as well as being able to dissect how the timing of locomotion (relative to whisker stimulation) impacted evoked haemodynamic responses. For the purpose of this study we were interested in how locomotion alters sensory-evoked haemodynamic responses, so for the 2s spontaneous dataset, time series analysis was not conducted (Figs 2-3), but spatial maps were generated (Fig 4).

## Results

### Locomotion alters the sensory-evoked hemodynamic response

Animals received a thinned cranial window surgery to allow 2D-OIS to measure changes in cerebral haemodynamics – specifically changes in oxygenated haemoglobin (HbO), deoxygenated haemoglobin (HbR) and total haemoglobin (HbT). During imaging, animals could move on a spherical treadmill whilst whiskers could be stimulated with a mechanical T-bar (Fig 1a). The treadmill was attached to an optical motion sensor, which allowed us to assess the impact that the amount and timing of locomotion (in isolation and also relative to whisker stimulation) had on the haemodynamic response. Twenty-one individual recording sessions, each with 59 trials, made up the 2s whisker stimulation data set. For each session, trials were ranked by voluntary locomotion occurring at different time points relative to the whisker stimulation. Evoked haemodynamic time series were generated from the top and bottom 10% of locomotion-ranked trials, corresponding to trials in which the most and least locomotion occurred, and were averaged across the 21 sessions.

**Figure 1.**
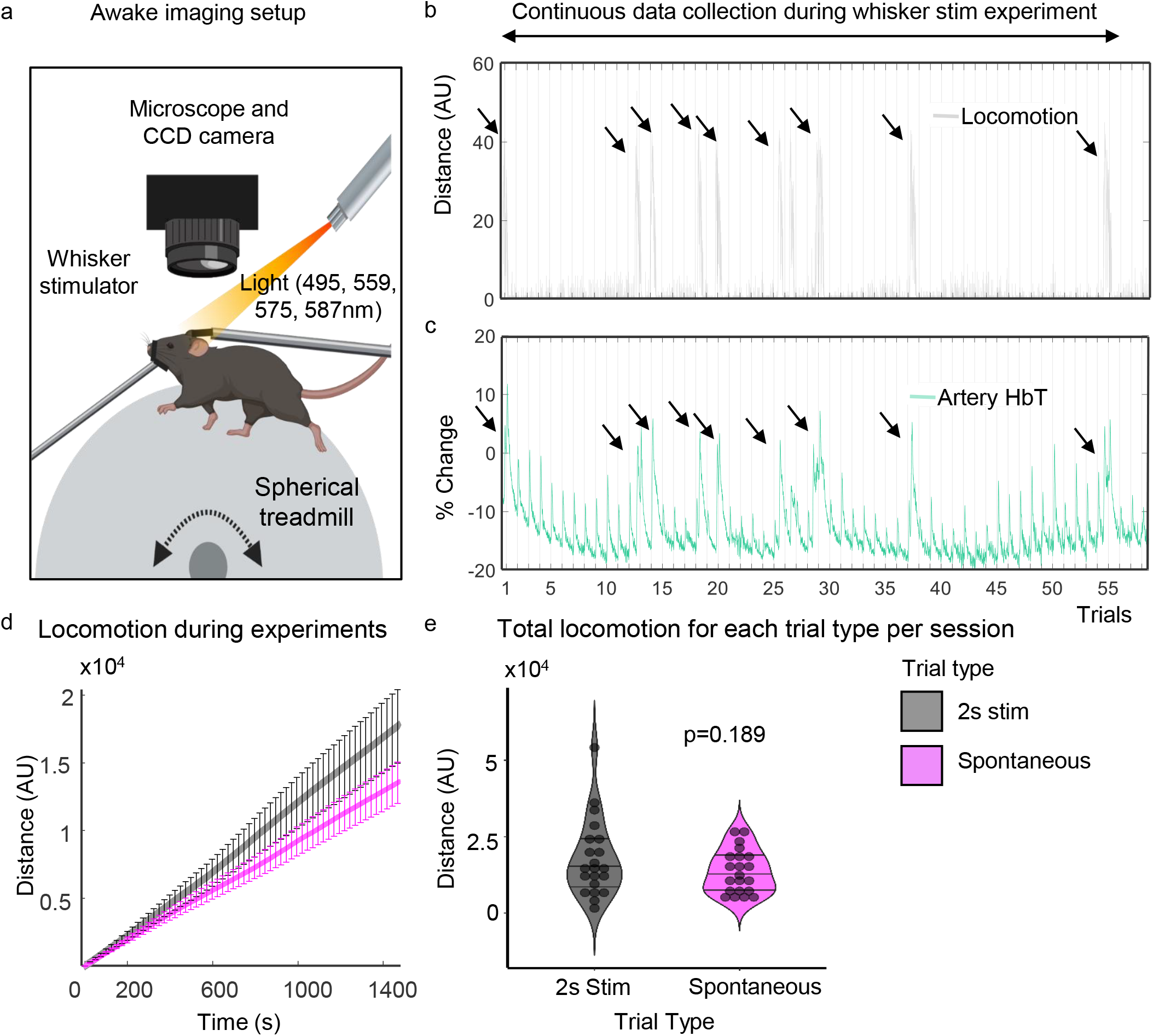
Awake imaging experimental set up. **(a)** Animals were head fixed and could run on a spherical treadmill. Locomotion data was collected using an optical motion sensor (attached to the ball – not shown). Light (495, 559, 575, 587 nm) was shone onto the thinned cranial window, and haemodynamic responses were collected during 2s whisker stimulation trials and during spontaneous (no whisker stimulation) trials. **(b)** and **(c)** show representative plots of the distance travelled (grey) and percentage change in HbT (green, taken from an artery within the whisker ROI (see Figure 4)) across the 2s whisker experiment (continuous data recording of the 59 whisker-stimulation trials taken from one representative animal/session, black lines on × axis mark individual 25s stimulation trials) respectively. Black arrows show large bouts of spontaneous walking. **(d)** shows total distance travelled during whisker stimulation trials (black/grey) compared to spontaneous trials (pink) averaged for all 21 sessions. Error bars represent +/-SEM. **(e)** shows a violin plot with individual points to show the distance travelled during each session for the different trial types (2s whisker stimulation and spontaneous trials with no whisker stimulation) (Sign test p=0.189). Black lines on violin plot represent interquartile range and median. Awake imaging experimental figure **(a)** created with BioRender.com

A representative locomotion and HbT response is shown in Figure 1b and c. Throughout the stimulations (marked on the x-axis), HbT responses to individual whisker stimulation trials can be observed. Large increases in HbT can be seen that coincide with spontaneous walking events (black arrowheads).

First, we checked whether the presence of whisker stimulation changes the amount of locomotion (Fig 1d and 1e). A sign test with continuity correction revealed there was no statistically significant difference in the median distance travelled (Median ± SEM: -1517 ± 2065) in experiments with a 2s whisker stimulation (Median ± SEM: 14894 ± 2732), compared to experiments without whisker stimulation (Median ± SEM: 12360 ± 1532, z = 1.309, p = 0.189).

We then looked at whether locomotion alters the evoked haemodynamic response. To do this we examined the effects of locomotion across the entire 25s trial period for the top and bottom 10% of trials ranked by locomotion. We examined how the greatest amount of locomotion influenced the evoked haemodynamic response as compared to the evoked haemodynamic response when the animals moved the least (Figs 2a, b and c). Sign tests with continuity correction revealed there were no statistically significant median differences in HbT peak (during a 2s whisker stimulation) (Median ± SEM: 0.003 ± .007) and HbO peak (Median ± SEM: -0.0004 ± .008) during trials with the greatest locomotion (Median ± SEM: HbT: 1.038 ± .007, HbO: 1.057 ± .009), as compared to when a stimulation occurred during trials with the least locomotion (Median ± SEM: HbT: 1.028 ± .003, z = -0.873, p=0.383, HbO: 1.049 ± .005, z = 0.000, p=1.000) (Figs 2b-e). Additionally, a Wilcoxon signed ranks test revealed no significant median difference in HbR peak (during a 2s whisker stimulation) (Median ± SEM: -0.0004 ± .005) during trials with the greatest locomotion (Median ± SEM: 0.948 ± .006), as compared to when a stimulation occurred during trials with the least locomotion (Median ± SEM: 0.945 ± .005, z = -0.330, p=0.741) (Figs 2b, c and f).

**Figure 2.**
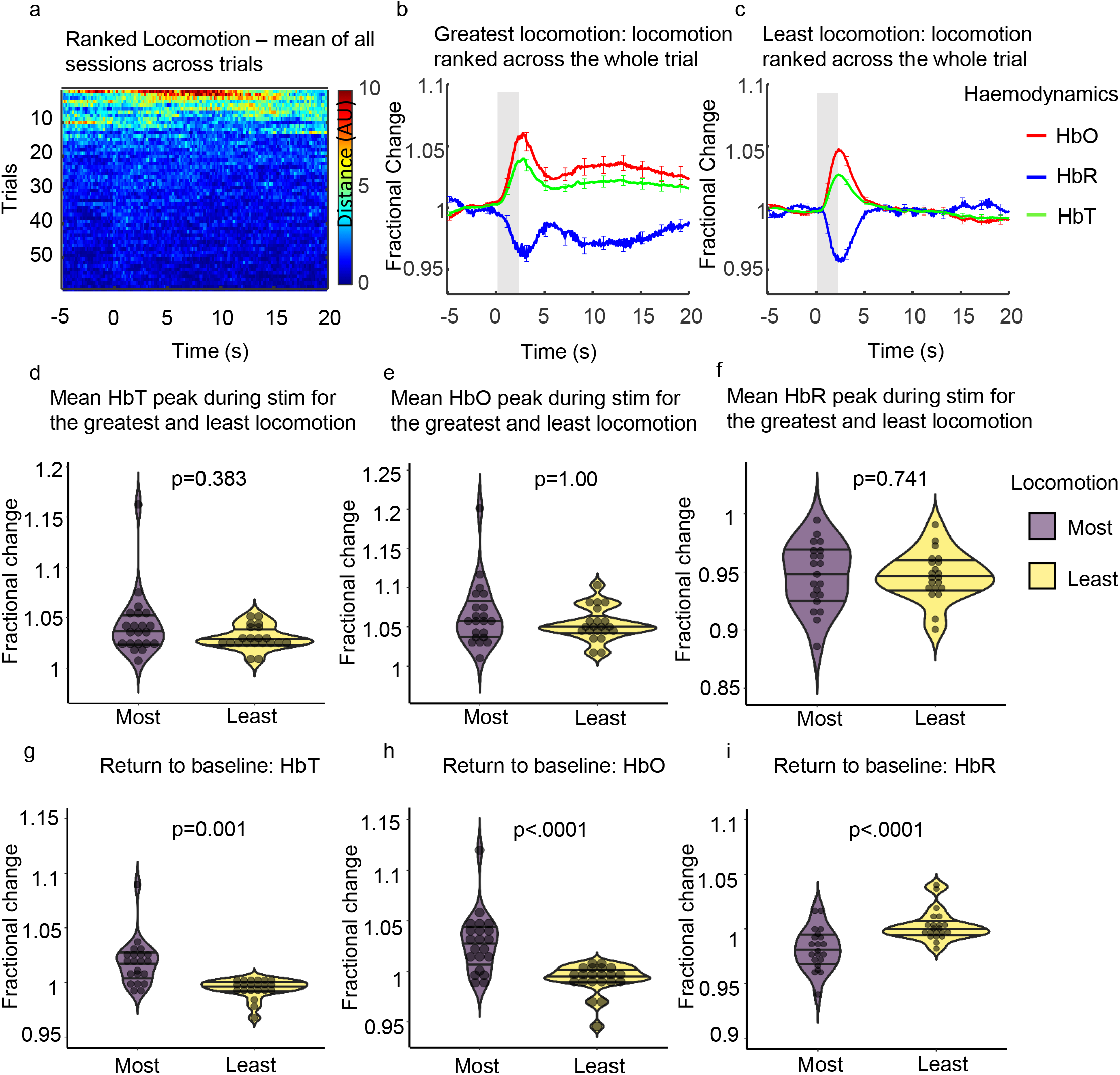
Mean sensory evoked haemodynamic responses in the whisker ROI for trials ranked with the greatest and least locomotion across the entire 25s trial. **(a)** Heat map showing locomotion traces for the 59 whisker-stimulation trials ranked by locomotion (across the whole 25s trial), **(b)** and **(c)** show mean fractional changes from baseline in HbO, HbR and HbT during a 2s whisker stimulation (grey shaded bar) when locomotion was ranked across the whole trial (between -5 to 20 seconds). ‘Greatest locomotion’ (left) represents the top 10% of these ranked trials, which correspond to trials with the most locomotion during the entire 25s trial recording (−5-20s) (21 sessions from 4 animals; per session n = 6 top trials (an average of the top ranked trials was taken for each session)). ‘Least locomotion’ (right) represents the bottom 10% of ranked trials, which correspond to trials in which the least amount of locomotion occurred during the 25s recording (21 sessions from 4 animals; per session n = 6 bottom trials (an average of the bottom ranked trials was taken for each session). Error bars represent mean +/-SEM between the total 126 trials. **(d), (e)** and **(f)** show mean sensory-evoked peak values for HbT, HbO and HbR respectively, for trials in which the most and least locomotion occurred when locomotion was ranked across the entire trial (25s). Violin plots show individual mean values overlaid. Black horizontal lines indicate interquartile range and median. P values are from Sign tests for HbT and HbO and from Wilcoxon Signed Ranks test for HbR. **(g), (h)** and **(i)** show the mean return to baseline values (mean values taken between 15-20s) for HbT, HbO and HbR respectively, for trials in which the most and least locomotion occurred when locomotion was ranked across the entire trial (25s). P values from Sign tests for HbT and HbR and Wilcoxon Signed ranks test for HbO. Black horizontal lines on violin plots indicate interquartile range and median.

However, in trials where the greatest locomotion occurred (Fig 2b) a slower return to baseline for HbT, HbO and HbR was observed (as compared to trials where the least locomotion occurred). We took mean values of HbT, HbO and HbR between 15-20s to assess the return to baseline differences across the two behaviours. Sign tests with continuity correction revealed a statistically significant median increase in HbT (Median ± SEM: 0.024 ± .005) and a median decrease in HbR (Median ± SEM: -0.021 ± .005) at the end of the 25s stimulation period during trials with the greatest locomotion (Median ± SEM: HbT: 1.019 ± .005, HbR: 0.980 ± .004) compared to trials where the least locomotion occurred (Median ± SEM: HbT: 0.997 ± .002, z = -3.055, p=.001, HbR: 0.999 ± .003, z = 3.491, p<.0001). A Wilcoxon signed ranks test revealed a statistically significant median increase in HbO (Median ± SEM: 0.032 ± .007) at the end of the 25s stimulation period during trials with the greatest locomotion (Median ± SEM: 1.030 ± .006) compared to trials where the least locomotion occurred (Median ± SEM: 0.994 ± .003, z = -3.702, p<.0001) (Figs 2b, c, g, h and i).

### The timing of locomotion (relative to whisker stimulation) impacts the sensory-evoked haemodynamic response

As we have shown that locomotion across the whole trial can alter the return to baseline of the sensory-evoked haemodynamic response we wanted to investigate in more detail how the timing of locomotion (relative to whisker stimulation) impacted the sensory-evoked response.

To do this, trials taken during the 2s whisker stimulation experiment were ranked by the amount of voluntary locomotion occurring across different 5s time windows (pre-stim: -5-0s, mid-stim: 0-5s, post-stim: 5-10s, 10-15s, 15-20s; Fig 3, Column 1). Evoked haemodynamic time series were created from the top and bottom 10% of ranked trials, these top and bottom 10% of ranked trials were averaged across sessions and corresponded to trials in which the most and least locomotion occurred (21 sessions; n = 6 top & n = 6 bottom per session (an average of the top and bottom ranked trials was taken for each session and used in the visualisation/analysis)) during the different 5s time windows (Fig 3). All mean peak values were taken between 0-5s and are referred to as occurring during the whisker stimulation.

**Figure 3.**
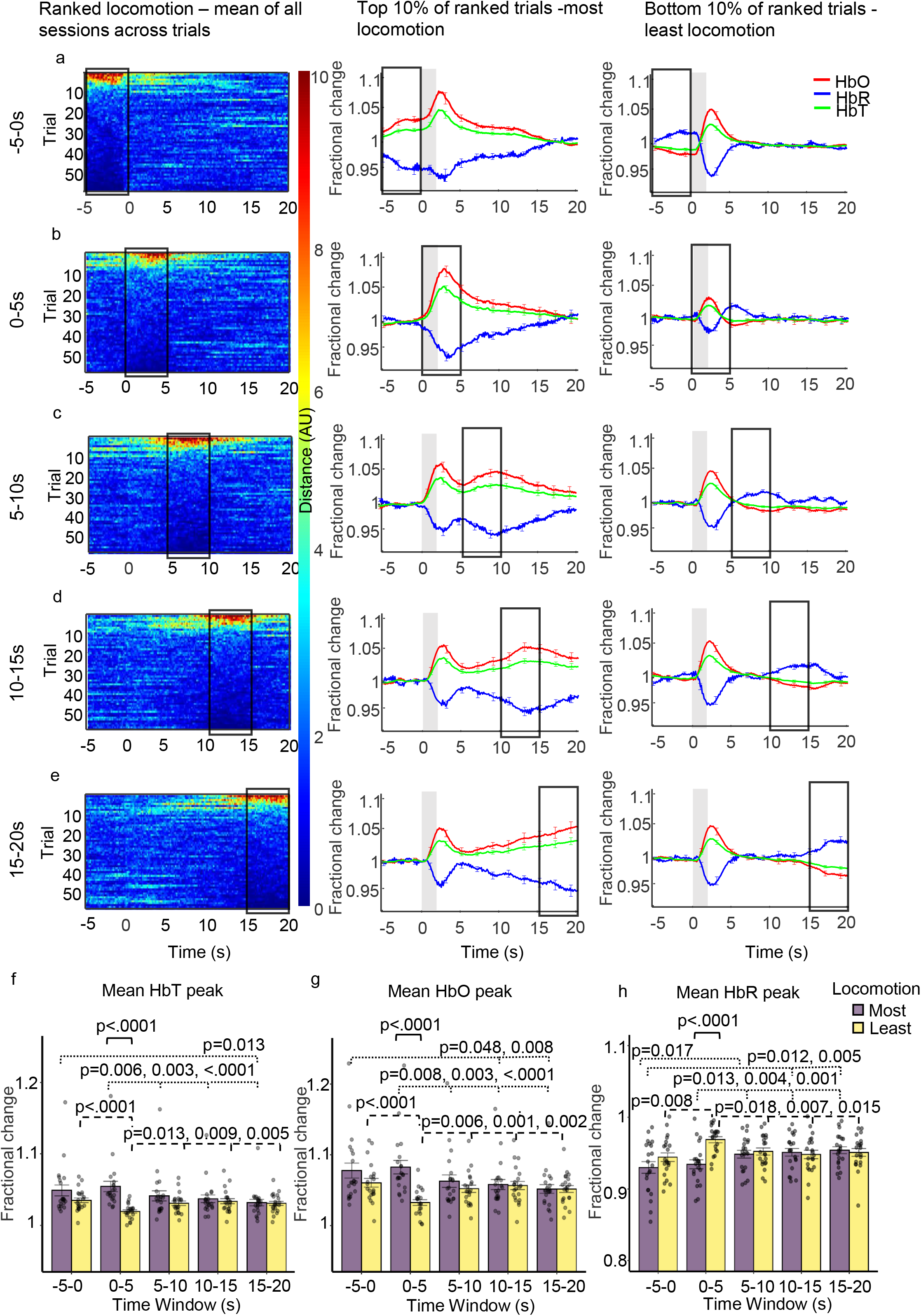
Mean sensory-evoked haemodynamic responses during trials where the most and least locomotion occurred with locomotion ranked at different time windows throughout the 25s trial. Whisker stimulation occurs between 0-2s (grey bar in centre and right columns). **Column one:** heat maps showing locomotion traces for the 59 whisker-stimulation trials, with locomotion ranked at different 5s time windows during the 25s trial (each ranked trial was averaged across 21 sessions/4 animals). The different 5s windows during the 25s trial where locomotion was ranked are: before (**a:** -5-0s), during (**b:** 0-5s) and after whisker stimulation (**c:** 5-10s, **d:** 10-15s, **e:** 15-20s). Trials were ranked according to locomotion in these 5s periods and presented in descending order. Colour bar indicates amount of locomotion, red pixels indicate more locomotion and dark blue indicate less locomotion. **Column two:** mean fractional changes from baseline in stimulation-dependent HbT, HbO and HbR taken from the top 10% of ranked locomotion trials across the different 5s time windows (21 sessions/4 animals; n=6 top per session (mean of top trials taken for each session, mean of all sessions used in the visualisation/analysis). **Column three:** mean fractional changes in stimulation-dependent HbT, HbO and HbR for the bottom 10% of locomotion trials ranked across the 5s time windows throughout the trial (21 sessions/4 animals; n=6 bottom per session (mean of bottom trials taken for each session, mean of all sessions used in the visualisation/analysis). Black boxes indicate the 5s time window locomotion was ranked. Data show mean across the total 126 trials +/-SEM. **(f), (g)** and **(h)** show mean +/-SEM between groups and individual mean peak values per session for HbT, HbO and HbR. Two-way repeated measures ANOVA’s were completed for HbT, HbO and HbR. Significant interactions were found and simple effects run (time and locomotion) for each of the haemodynamic measures. P values from pairwise comparisons (Bonferroni correction) are reported. Black solid brackets indicate comparison between most and least locomotion, dotted brackets reveal comparisons for most locomotion across different time windows and dashed brackets show comparisons for least locomotion across different time windows.

Three two-way repeated measures ANOVAs for HbT, HbO and HbR respectively, revealed that there was a significant interaction between the amount of locomotion (factors: most & least) and the time at which locomotion was ranked (factors: -5-0s, 0-5s, 5-10s, 10-15s, 15-20s) on the peak of the haemodynamic response to the 2s whisker stimulation (peak occurring between 0-5 seconds):-HbT: *F*(2.58, 51.57) = 13.35, *p*<.0001, ε = .645; HbO: *F*(2.89, 57.88) = 13.32, *p*<.0001, ε = .723; HbR: *F*(2.53, 50.52) = 8.712, *p* <.0001, ε = .632), indicating that the effect of locomotion was dependent on the timing of locomotion.

To dissect how the timing of locomotion during the trial impacted the sensory-evoked haemodynamic response (HbT, HbO or HbR peak detected between 0-5s), simple main effects were run to assess how ranked-locomotion during the five different time windows impacted the sensory-evoked haemodynamic response.

#### Most Locomotion trials

Simple main effects revealed that for the trials in which the most locomotion occurred (Fig 3, Column 2), the time at which locomotion was ranked had a significant effect on the mean peak of HbT (*F*(2.52, 50.41) = 12.99, *p* <.0001, ε = .630), HbO *(F*(2.66, 53.19) = 12.79, *p* = <.0001 ε = .665), and HbR (*F*(2.69, 53.78) = 11.50, *p* <.0001, ε = .672) during the 2s whisker stimulation. Pairwise comparisons with a Bonferroni correction revealed significant differences when locomotion occurred before (−5-0s) and during the stimulation (0-5s), as discussed in detail below (Table 1).

**Table 1:** Comparisons between 5s time windows for HbT, HbO and HbR peaks during whisker stimulation for most and least locomotion conditions. Mean diff refers to mean difference between the time window in which locomotion was ranked, compared with the time window compared to. Pairwise comparisons with a Bonferonni correction reported. SEM is standard error of the mean. * indicates a significant difference between the means.

| Time Window |  | Haem Measure | Most Locomotion |  |  | Least Locomotion |  |  |
| --- | --- | --- | --- | --- | --- | --- | --- | --- |
| Locomotion ranked | Compared to |  | Mean diff | SEM | p | Mean diff | SEM | p |
| -5-0 | 0-5 | HbT | -.006 | .003 | 1.000 | .016 | .003 | .000* |
|  |  | HbO | -.005 | .006 | 1.000 | .028 | .005 | .000* |
|  |  | HbR | -.004 | .006 | 1.000 | -.025 | .006 | .008* |
|  | 5-10 | HbT | .008 | .003 | .194 | .004 | .002 | .769 |
|  |  | HbO | .015 | .005 | .065 | .008 | .004 | .356 |
|  |  | HbR | -.018 | .005 | .017* | -.008 | .004 | .546 |
|  | 10-15 | HbT | .012 | .004 | .100 | .001 | .003 | 1.000 |
|  |  | HbO | .020 | .006 | .048* | .004 | .005 | 1.000 |
|  |  | HbR | -.021 | .006 | .012* | -.004 | .005 | 1.000 |
|  | 15-20 | HbT | .017 | .005 | .013* | .004 | .003 | 1.000 |
|  |  | HbO | .026 | .007 | .008* | .009 | .006 | 1.000 |
|  |  | HbR | -.024 | .006 | .005* | -.007 | .007 | 1.000 |
| 0-5 | 5-10 | HbT | .013 | .003 | .006* | -.011 | .003 | .013* |
|  |  | HbO | .020 | .005 | .008* | -.020 | .005 | .006* |
|  |  | HbR | -.014 | .004 | .013* | .017 | .005 | .018* |
|  | 10-15 | HbT | .017 | .004 | .003* | -.014 | .004 | .009* |
|  |  | HbO | .025 | .006 | .003* | -.024 | .005 | .001* |
|  |  | HbR | -.017 | .004 | .004* | .021 | .005 | .007* |
|  | 15-20 | HbT | .023 | .004 | .000* | -.011 | .003 | .005* |
|  |  | HbO | .031 | .006 | .000* | -.019 | .004 | .002* |
|  |  | HbR | -.020 | .004 | .001* | .018 | .005 | .015* |
| 5-10 | 10-15 | HbT | .004 | .002 | .933 | -.003 | .002 | 1.000 |
|  |  | HbO | .005 | .003 | 1.000 | -.005 | .003 | 1.000 |
|  |  | HbR | -.003 | .003 | 1.000 | .004 | .004 | 1.000 |
|  | 15-20 | HbT | .010 | .004 | .158 | .000 | .003 | 1.000 |
|  |  | HbO | .011 | .005 | .346 | .000 | .005 | 1.000 |
|  |  | HbR | -.006 | .004 | 1.000 | .001 | .005 | 1.000 |
| 10-15 | 15-20 | HbT | .005 | .002 | .232 | .003 | .003 | 1.000 |
|  |  | HbO | .006 | .003 | .344 | .005 | .005 | 1.000 |
|  |  | HbR | -.003 | .003 | 1.000 | -.003 | .005 | 1.000 |

There were notable differences in the mean peaks of HbT, HbO and HbR during the 2s whisker stimulation (as assessed by pairwise comparisons with a Bonferroni correction). When locomotion was ranked before the stimulation (−5 to 0s), mean HbO peak during the 2s stimulation was greater than when locomotion was ranked at 10-15 and 15-20s and mean HbT peak during the 2s stimulation was greater than when locomotion was ranked at 15-20s(Fig 3f-g). Additionally, when locomotion was ranked before the stimulation (−5-0) mean HbR peak during the 2s stimulation was less than when locomotion was ranked at all time windows after the stimulation (5-10s, 10-15s and 15-20s) (Fig 3h) - indicating a larger HbR washout occurs when locomotion is ranked before the whisker stimulation as compared to after the whisker stimulation (Table 1).

Additionally, when locomotion was ranked at the stimulation onset (0-5s) mean HbT and HbO peak responses during the 2s stimulation were greater than when locomotion was ranked at all time windows after the stimulation (Fig 3f-g). For HbR when locomotion was ranked at stimulation onset (0-5s) mean HbR peak during the 2s whisker stimulation was less than when locomotion was ranked at all time windows after the stimulation (Fig 3h) - indicating a larger HbR washout when locomotion was ranked at stimulation onset as compared to when it occurred after the whisker stimulation (Table 1).

#### Least Locomotion trials

Simple main effects revealed that for the trials in which the least locomotion occurred (Fig 3, Column 3), the time at which locomotion was ranked also had a significant effect on the mean peak of HbT (*F*(2.94, 58.79) = 8.93, *p* <.0001, ε = .735), HbO (*F*(4, 80) = 9.94, *p* <.0001) & HbR (*F*(4, 80) = 6.83, *p* <.0001) during the 2s whisker stimulation. Pairwise comparisons with a Bonferroni correction revealed that significant differences were observed when locomotion was ranked before the stimulation (−5-0s) and at the stimulation onset (0-5s) vs the post-stimulation ranked locomotion conditions (5-10s, 10-15s, 15-20s).

When locomotion was ranked before the stimulation (−5-0s), the mean HbT and HbO peak responses were greater than when locomotion was ranked at stimulation onset (0-5s) (Fig 3f-g). Additionally, mean HbR peak when locomotion was ranked before the stimulation (−5-0s) was less than when locomotion was ranked at stimulation onset (0-5s) (Fig 3h) – indicating a larger HbR washout when locomotion was ranked before the stimulation, as compared to when ranked during the stimulation (Table 1).

When locomotion was ranked at the stimulation onset (0-5s) mean HbT and mean HbO peaks during the 2s whisker stimulation were less than when locomotion was ranked at all time windows after the stimulation (Fig 3f and 3g). Whereas mean HbR peak during whisker stimulation when locomotion was ranked at stimulation onset (0-5s) was greater than when locomotion was ranked at all time windows after the stimulation (Fig 3h) – indicating a smaller HbR washout when locomotion was ranked at stimulation onset compared to when ranked after the stimulation (Table 1).

### The amount of locomotion impacts the sensory-evoked haemodynamic response only when locomotion is ranked at specific time windows

Having previously shown that there is a significant interaction between the amount of locomotion (factors: most & least) and the time at which locomotion was ranked (factors: -5-0s, 0-5s, 5-10s, 10-15s, 15-20s) on the peak of the haemodynamic response to the 2s whisker stimulation (peak occurring between 0-5 seconds), as well as highlighting at which time windows ranked-locomotion impacted the sensory-evoked haemodynamic response, we now wanted to reveal how the amount of locomotion at these five time windows impacted the sensory-evoked response.

Simple main effects with a Bonferroni correction revealed that mean whisker stimulation-evoked HbT peak was greater for trials in which the most locomotion occurred as compared to trials in which the least locomotion occurred when locomotion was ranked at stimulation onset (0-5s; *F*(1,20) = 19.68, *p*<.0001, mean ± SEM: 1.054 ± .007 vs 1.020 ± .002) (Figs 3a, b, Columns 2 and 3). No significant differences were found when comparing the effect of locomotion on the mean whisker stimulation-evoked HbT peak when locomotion was ranked before the stimulation (−5-0) and at 5-10s, 10-15s and 15-20s. This indicates that the amount of locomotion only effects the HbT element of the evoked-haemodynamic response when locomotion occurs during the stimulation.

Simple main effects with a Bonferroni correction indicated that mean whisker stimulation-evoked HbO peak was also greater for trials in which the most locomotion occurred as compared to trials in which the least locomotion occurred when locomotion was ranked at stimulation onset (0-5s; *F*(1,20) = 24.83, *p*<.0001, mean ± SEM: 1.083 ± .009 vs 1.033 ± .004) (Figs 3a, b, Columns 2 and 3). No significant differences were found when comparing the effect of locomotion on the mean whisker stimulation-evoked HbO peak when locomotion was ranked before the stimulation (−5-0) and when ranked at 5-10s, 10-15s and 15-20s. Indicating that the amount of locomotion only effects the HbO element of the evoked-haemodynamic response when locomotion occurs during the stimulation.

Simple main effects with a Bonferroni correction revealed that mean whisker stimulation-evoked HbR peak was less for trials in which the most locomotion occurred as compared to trials in which the least locomotion occurred when locomotion was ranked at stimulation onset (0-5s; *F*(1,20) = 38.75, *p*<.0001, mean ± SEM: 0.934 ± .006 vs 0.969 ±.004) – indicating a larger HbR washout when the animal moved more (Figs 3a, b, Columns 2 and 3). No significant differences were found when comparing the effect of locomotion on the mean whisker stimulation-evoked HbR peak for trials in which locomotion was ranked before the stimulation (−5-0) and when ranked at 5-10s, 10-15s and 15-20s. This indicates that the amount of locomotion only effects the HbR element of the evoked-haemodynamic response when locomotion occurs during the stimulation.

### Locomotion impacts the spatial spread of HbT across the surface vasculature

Representative spatial maps from each animal show HbT activation - revealing fractional change in HbT within the surface vasculature, during locomotion occurring with and without a 2s whisker stimulation (activation between 0-5s) (Fig 4). Red pixels indicate increased activation and blue pixels decreased activation. Figure 4 (Column 2) was generated from spontaneous trials (with no stimulation). Column 2 shows that during locomotion, a more global activation can be observed which is not restricted to the whisker region alone (red ROIs, Column 1) as per when a whisker-stimulation occurs concurrently with limited locomotion (see Column 4, ‘least’ locomotion). Figure 4 (Columns 3 & 4) shows representative spatial maps for each animal for the trials in which the most and least locomotion occurred during the 2s whisker stimulation. Increased activation can be observed within the whisker region (red outline, Fig 4, Column 1) for trials in which the most locomotion occurred during the 2s whisker stimulation (Fig 4, Column 3). Increased activation within the whisker region can also be observed in trials where animals moved the least during a whisker stimulation. A decrease in activation (blue pixels in a region adjacent to the whisker region within red outline) can also be seen, which suggests a reduction in HbT within a region anterior to the whisker area (Fig 4, Column 4).

**Figure 4.**
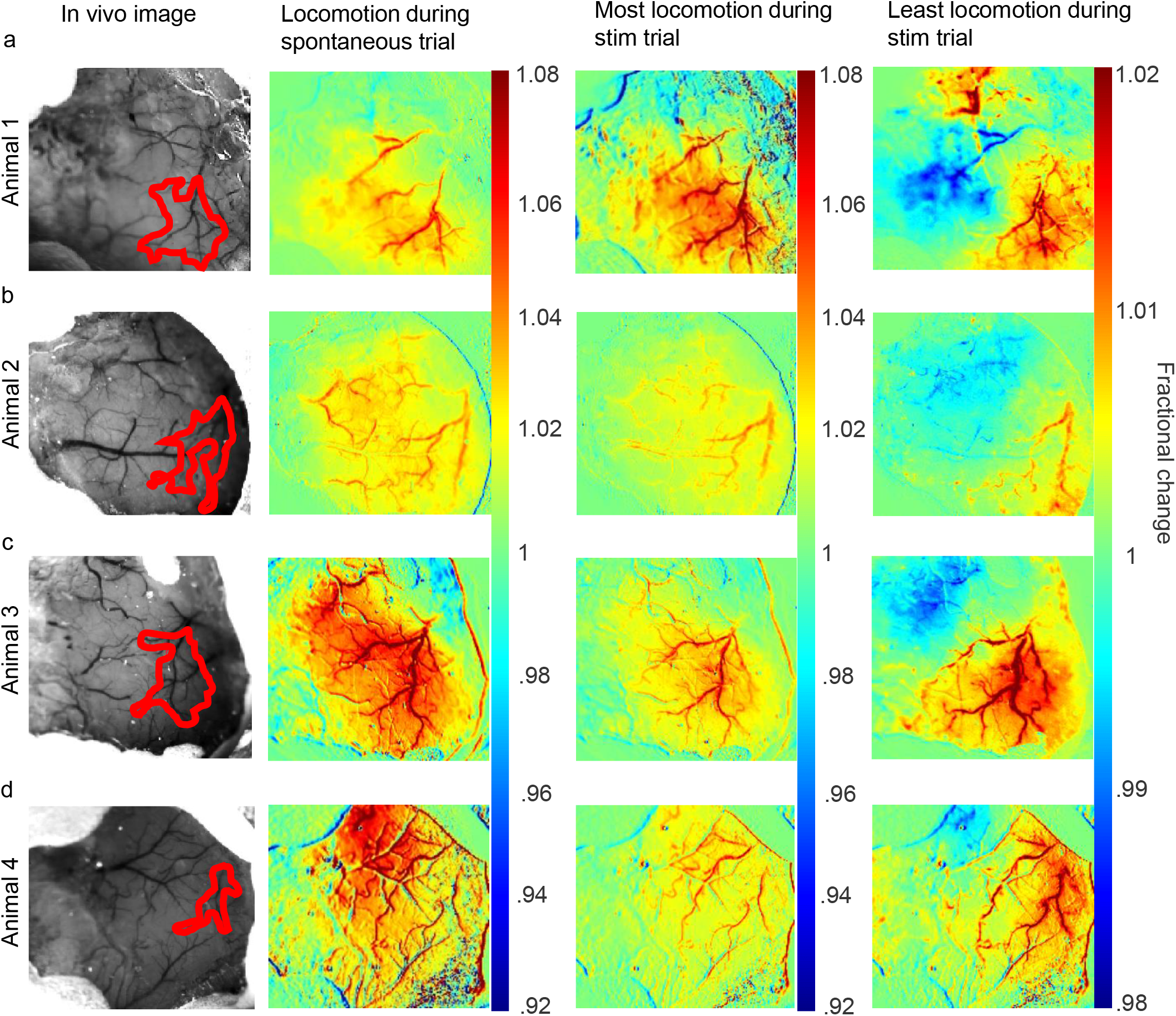
Representative HbT spatial maps during locomotion. Spatial maps from each animal included in the analysis **(a-d)** showing the surface vasculature in the somatosensory cortex as recorded during locomotion alone (left centre, spontaneous recordings), and during trials with the most (right centre) and least (right) locomotion occurring during the 2s whisker stimulation (from the 0-5s time window, see Fig 3b). **Column one:** in vivo images of the thinned cranial window with the automatically generated whisker region highlighted in red. **Column two:** spatial maps showing fractional changes in HbT generated from spontaneous trials (with no whisker stimulation) during 25s bouts of continuous locomotion. **Column three & four:** HbT spatial maps of trials in which the most (right centre) and least (right) locomotion occurred during a 2s whisker stimulation. This map reveals the spatial location of the whisker region (red pixels, which corresponds to the automatically generated whisker region in red ROI of Column 1), as well as revealing an area with a decrease in fractional change of HbT (blue pixels). Colour bar represents fractional change in HbT, with red indicating an increase in fractional change and blue indicating a decrease. Column 1, **(b)** ((in vivo image animal 2) also used in Sharp et al., 2015^20^, see Figure 4B).

## Discussion

The present study measured spontaneous (Fig 4) and sensory-evoked (Figs 2-4) haemodynamic responses from the cerebral cortex in head-fixed, awake mice, whilst locomotion was concurrently monitored. The novel aspect of our approach was to investigate the impact of the amount and timing of locomotion events on sensory-evoked haemodynamic responses. Our experiments revealed that sensory-evoked haemodynamic responses are altered by the presence of locomotion, which was dependent on the timing (relative to whisker stimulation) that the locomotion occurred. Our findings suggest there is a relationship between the time at which locomotion occurs (in 5s time windows relative to the whisker stimulation) and the amount of summed locomotion, and that this affects the evoked haemodynamic response, with locomotion appearing to have the largest effects when it occurred before the stimulation (−5-0s) and during (0-5s) the stimulation. We therefore suggest that it is especially important to monitor locomotion in awake imaging experiments in which haemodynamics are being assessed/compared between groups (e.g., disease vs healthy subjects).

Previous studies have shown that certain behaviours, including body movements and whisking can enhance CBV in awake, head-fixed mice^28^. It has also been reported that locomotion, in the absence of sensory stimulation, increases cortical CBV^23^, and rapidly dilates arteries^25^. However, the above studies did not explore how locomotion may specifically affect sensory-evoked haemodynamic responses, making our findings novel.

Of the few papers that have investigated the impact of locomotion on evoked haemodynamic responses, Tran et al., (2018)^24^ explored whether the behaviour of an animal during whisker stimulation had an effect on penetrating arteriole dilation. They showed that the behaviour of the animal did not alter the peak amplitude of arteriole dilation. In comparison, our data suggests that the animal’s behaviour does impact the peak amplitude of arteriole dilation. Our opposing findings could be explained by the different methods used in the two studies. Tran et al. used two-photon microscopy to investigate haemodynamic responses, focusing on penetrating vessel dilation. In contrast, our study used 2D-OIS to measure changes in blood oxygenation from the surface of the cerebral cortex. It is conceivable that locomotion and the time at which locomotion occurs relative to whisker stimulation may have a differing impact on evoked haemodynamic responses depending on the type and location of blood vessels investigated. In support of this explanation, Gao et al., (2015)^27^ found that locomotion impacted surface vessels to a greater extent than penetrating vessels, with locomotion leading to surface vessel dilations that were almost three times the size of intracortical vessel dilation. However, the study did not investigate how locomotion affected sensory-evoked haemodynamic responses. Future studies using two-photon microscopy are still warranted^29^ to explore how sensory-evoked haemodynamic responses are impacted by locomotion, as well as to assess if our results can be replicated using other methods.

Our study was not without limitations, the behavioural set up only monitored locomotion behaviours and did not monitor whisking behaviours or pupil dilations in the animals. It has been reported that whisking occurs when an animal moves^30^ Therefore, there may also be a relationship between locomotion, whisking and sensory-evoked haemodynamic responses.

Future studies may benefit from monitoring both locomotion and whisking^24^ to assess if there are interactions between these behaviours and the impact this may have on the evoked haemodynamic response – as voluntary whisking has been reported to increase CBV^28^. However, the purpose of our paper was to focus on the effects of locomotion on sensory-evoked haemodynamic responses.

Additionally, neural activity was not measured during the study, it would have been informative to observe how neural activity was affected by the amount and timing of locomotion. Other studies have recorded neural activity alongside CBV during voluntary locomotion and found that voluntary locomotion does indeed increase neural activity^23,31^. Measuring simultaneous haemodynamics in awake animals combined with genetically encoded calcium indicators (such as GCAMP6)^7^ to measure spontaneous and evoked neuronal activity will provide additional information on how brain activity is modulated by the interaction of locomotion and sensory stimulation.

Our paper demonstrates the importance of monitoring behaviour – especially locomotion - during awake haemodynamic imaging. As our study shows that the amount and timing of locomotion (relative to whisker stimulation) can impact the amplitude of an evoked haemodynamic response we suggest that, where possible, groups should monitor locomotion in their awake imaging experiments – particularly when using sensory stimulation. If locomotion behaviours cannot be monitored, other methods could be used to limit locomotion behaviours, such as training animals to remain stationary^32^. Monitoring locomotion is especially important to consider when comparing different disease groups, in which locomotion may differ^33^ – if locomotion behaviour is not monitored (or excluded), confounded conclusions could potentially be made.

## Methods

### Animals

Adult (3-12m; 24-40g) female C57/BL6J mice (n = 4) were used in the experiment. Food and water were available ad-libitum and mice were housed on a 12hr dark/light cycle. All animal procedures were approved by the UK Home Office and in agreement with the guidelines and scientific regulations of the Animals (Scientific Procedures) Act 1986 with additional approval received from the University of Sheffield licensing committee and ethical review board.

### Surgery

Induction of anaesthesia was achieved with a combination of fentanyl-fluanisone (Hypnorm, Vetapharm Ltd), midazolam (Hypnovel, Roche Ltd) and sterile water in the ratio 1:1:2 (1ml/kg i.p). Surgical anaesthetic plane was maintained using isoflurane (0.25-0.8%) in 100% oxygen. Body temperature was monitored and maintained throughout surgery via a rectal thermometer and a homeothermic blanket respectively (Harvard Apparatus). Eyes were protected using Viscotears (Novartis). A scalpel was used to shave the head prior to the mouse being positioned in a stereotaxic frame (Kopf Instruments). Iodine was applied to the scalp and the scalp was removed. Using a dental drill, the bone covering the right somatosensory cortex was thinned to translucency to create the thinned optical window (∼3mm^2^). Cyanoacrylate glue was thinly applied across the window to strengthen the window and reduce optical specularities. Dental cement (Superbond C & B; Sun Medical) was applied to the bone on the contralateral side of the cranial window and a well was built up around the window to allow for a metal head plate to be attached for chronic imaging. Following surgery, mice were housed individually and given at least one week to recover before any imaging commenced.

### Awake imaging

Prior to imaging, mice were gradually habituated to the experimenter, imaging room, spherical treadmill and head-fixation. To achieve this, training sessions were completed with a reward at the end of each session (toffee popcorn, Sunkist). The first session lasted approximately 10 minutes. The experimenter handled the mice and allowed the mice to explore the spherical treadmill without head fixation. The second session was a repeat of the first session. Session three involved head-fixing the mice for approximately 10 minutes whilst the lights were on. This was followed by ∼20 minutes with the lights off. Session three was repeated daily until mice learned how to move on the spherical treadmill and displayed grooming behaviours (approximately 2-3 sessions). The whisker stimulator was introduced during the final two training sessions.

### Whisker Stimulation

Whiskers were mechanically stimulated using a plastic T-bar at 5Hz. Each experiment lasted 1475 seconds and comprised of 59 25s trials. During whisker stimulation trials whisker deflection lasted 2s, occurring every 25s. Spontaneous experiments were also conducted using the same timings as 2s whisker stimulation experiments, however the motor controlling the whisker stimulator was switched off, ensuring whiskers were not stimulated.

### Locomotion Data Collection and Analysis

Locomotion data was collected from a spherical treadmill with an optical motion sensor attached, to quantify locomotion. Locomotion data was analysed using in-house created scripts in MATLAB (MathWorks). The optical motion sensor recorded the movement of the treadmill and produced a file comprised of: locomotion data (a vector which showed the rotation of the treadmill, integers were used to quantify the displacement of the treadmill, with stationary periods reflected by 0, the quicker the spherical treadmill moved, the higher the integer; plotted as distance (arbitrary unit, AU)); the time vector (which allowed locomotion to be measured across time (s)); and the trigger points (these indicated the timing of the whisker stimulation, across trials, this enabled locomotion data to be matched with the timing of the haemodynamic data). To establish if locomotion did impact evoked-haemodynamic responses, 2s whisker stimulation trials were ranked by voluntary locomotion across the entire trial (25s) and across different 5s time windows within the stimulation period (−5-0s, 0-5s, 5-10s, 10-15s, 15-20s). For each session, evoked haemodynamic time series were created from the top and bottom 10% of ranked trials, these top and bottom 10% of ranked trials were averaged together across sessions and corresponded to trials in which the most and least locomotion occurred (21 sessions from 4 animals, n = 6 top & n = 6 bottom trials per session (an average of the top and bottom ranked trials was taken for each session and used in the visualisation/analysis)). In Figure 4 (Column 2), HbT spatial maps for spontaneous locomotion were created as followed. Locomotion events from spontaneous trials were selected and a spectroscopy file was created to assess how locomotion alone impacts the spatial spread of HbT within the surface vasculature

### 2D-Optical Imaging Spectroscopy (2D-OIS)

2D-OIS uses light to measure cortical haemodynamic signals by estimating concentration changes in oxygenated haemoglobin (HbO), deoxygenated haemoglobin (HbR) and total haemoglobin (HbT). In order to measure changes in cortical haemodynamics a Lambda DG-4 high-speed galvanometer (Sutter Instrument Company, USA) was used to illuminate the right somatosensory cortex with 4 wavelengths of light (495 ± 31nm, 559 ± 16nm, 575 ± 14nm and 587 ±nm). A Dalsa 1M60 CCD camera was used to capture remitted light at 184 × 184 pixels, at a 32 Hz frame rate, this provided a resolution of ∼75µm.

To produce 2D images of micromolar changes in HbO, HbR and HbT, spectral analysis (based on the path length scale algorithm (PLSA)) was conducted^34,35^. This algorithm uses a modified Beer Lambert Law, with a path-length correction factor and predicted absorption values of HbO, HbR and HbT. The relative concentration estimates of HbO, HbR and HbT were gathered from baseline values, whereby haemoglobin tissue concentration was estimated as 100 µM, with tissue saturation of oxygen estimated at 80%.

### Regions of Interest (ROI) overlying the whisker barrels from 2D spatial maps

MATLAB (MathWorks) was used to select ROI for time series analysis. Custom-made in-house scripts were used to select ROIs from the 2D spatial maps produced using 2D-OIS. The whisker ROI was selected using the HbT spatial map taken from the 2s whisker stimulation experiments; this was completed for each of the 21 sessions. Pixels were included in the ‘active’ region if they were > 1.5 × STD across the entire spatial map, hence the whisker ROI (red ROI, Fig 4, Column 1) was the area of cortex with the greatest haemodynamic response for HbT. The following time series analyses included in the study (Fig 2 & 3) were conducted for the whisker region.

### Statistical Analysis

Statistical tests were conducted in SPSS (v26) and figures were created in MATLAB and RStudio. P values of <0.05 were deemed to be significant. Outliers were assessed using box plots, with values greater than 1.5 box lengths from the edge of the box classified as outliers – outliers were kept in the data set. Normality was assessed using the Shapiro Wilk test. If outliers were observed and/or data was non-normal, non-parametric tests were used (if available).

For distance travelled calculations, the total distance (AU) from each session for experiments with and without whisker stimulation were used – with the total sum of distance travelled taken for each experiment during each of the 21 sessions. A sign test was used to assess if there was a statistically significant difference in distance travelled during experiments.

Non-parametric Sign tests (HbT and HbO) and the Wilcoxon signed ranks test (HbR) were used to assess if there were significant differences in HbO, HbR and HbT peaks during the 2s whisker stimulation when comparing trials with the greatest and least amount of locomotion (when locomotion was ranked across the entire 25s). The peak amplitude of HbT, HbO and HbR were computed as the time point with the greatest change in the concentration of haemoglobin from baseline^20^ between 0-5 seconds during the ranked trials where the most and least locomotion occurred.

Sign tests (HbT and HbR) and the Wilcoxon signed ranks test (HbO) were used to establish if there were significant differences in the return to baseline of HbT, HbO and HbR, when comparing trials in which the most and least locomotion occurred during a 2s whisker stimulation – mean HbT, HbO and HbR values were taken at the end of the 25s stimulation period (mean values between 15-20s).

Three, two-way repeated measures ANOVAs were completed to assess if there was an effect of the amount (factors: most & least) and timing (factors: -5-0s, 0-5s, 5-10s, 10-15s, 15-20s) of locomotion on the peak of the haemodynamic response (dependent variables: HbT, HbO, HbR) to the 2s whisker stimulation (peak occurring between 0-5 seconds). The presence of outliers was assessed using studentised residuals, where values greater than ±3 were deemed to be outliers. Outliers were observed and were kept in the data set. Normality was assessed by the Shapiro Wilk test, and sphericity was assessed using the Mauchly’s test of sphericity. For the two-way ANOVAs, a number of variables were not normally distributed (see Supplementary Statistics Table S9). If Mauchly’s sphericity was violated (p <.05) Greenhouse Geiser correction was used. The use of the Greenhouse Geiser correction can be observed if there is an epsilon (ε) value when reporting ANOVA results. As there is no non-parametric alternative for a two-way ANOVA if variables were not normally distributed and outliers were present, a two-way ANOVA was still completed, as ANOVAs are robust to slight deviations from normality. Data were not transformed as transforming the data results in difficulties comparing the means across different groups^36^. If an interaction effect was found, to assess the simple main effects, one-way ANOVAs were completed and pairwise comparisons with a Bonferroni correction were completed. Data are reported as means ± standard error of the mean (SEM), unless otherwise stated. Individual dots on violin plots and bar charts represent individual mean data points. Data was visualised as a bar plot when statistical tests compared the mean, whereas violin plots were used when statistical tests compared the median. Detailed statistical outputs can be found in the supplementary tables.

## Supporting information

Supplementary statistics tables

## Data availability

Data sets used/analysed in the current study are available in the DRYAD repository, https://doi.org/10.5061/dryad.v41ns1rxs

## Acknowledgements

BE would like to thank the Battelle – Dr Jeff Wadsworth studentship for funding her PhD [Grant number R/156208-14-1]. OS is funded by a British Heart Foundation (BHF) project grant (PG/13/55/30365). CH is funded by a Sir Henry Dale Fellowship jointly funded by the Wellcome Trust and the Royal Society. This research was funded in whole, or in part, by the Wellcome Trust [Grant number 105586/Z/14/Z]. For the purpose of Open Access, the author has applied a CC BY public copyright licence to any Author Accepted Manuscript version arising from this submission. We also would like to thank Michael Port for building and maintaining the whisker stimulation device and 2D-OIS imaging equipment. Awake imaging experimental figure created with BioRender.com.

## Author Information

### Contributions

B.E wrote the main manuscript. C.H and J.B conceived research ideas. K.S and P.S completed surgeries and imaging experiments. B.E, J.B and C.H analysed the data. L.B, K.S., B.E., C.H & J.B. wrote the code for analysis. K.S, L.L, O.S, C.H, and J.B edited and proofread the manuscript. B.E and K.S contributed equally. J.B and C.H contributed equally.

