## Supplementary statistics tables for "The effects of locomotion on sensory-evoked haemodynamic responses in the cortex of awake mice"

**Supplementary tables for statistics outputs**

All data values were included in the statistical analysis even if outliers were found as there was no scientific reason to remove them from the data set. Data used in the analysis can be found in the DRYAD repository, doi:10.5061/dryad.v41ns1rxs

Distance travelled

**Supplementary table S1: Shapiro Wilk test of normality for distanced travelled during spontaneous and whisker stimulation trials**

| Distance travelled | Statistic | df | p |
|---|---|---|---|
| Spontaneous vs Whisker stim | .691 | 21 | .000 |

*Normality was assessed using a difference variable comparing distanced travelled during spontaneous trials with distance travelled during whisker stimulation trials*

**Supplementary table S2: Sign test result for distance travelled during spontaneous and whisker stimulation trials**

| Type of trial | N | Mean | Median | SEM | z | p |
|---|---|---|---|---|---|---|
| Spontaneous | 21 | 13610 | 12360 | 1532 | - | - |
| 2s whisker stim | 21 | 17802 | 14894 | 2732 | - | - |
| Difference | 21 | -4192 | -1517 | 2065 | 1.309 | .189 |

Greatest vs Least locomotion

**Supplementary table S3: Shapiro Wilk test of normality for greatest vs least locomotion (where locomotion ranked across the entire 25s trial)**

| | Greatest Locomotion vs Least Locomotion | | |
|---|---|---|---|
| **Haemodynamic Measure** | **Statistic** | **df** | **p** |
| HbT | .568 | 21 | .000 |
| HbO | .695 | 21 | .000 |
| HbR | .923 | 21 | .100 |

**Supplementary table S4: Wilcoxon signed ranks result for HbR greatest vs least locomotion (where locomotion is ranked across the entire 25s trial)**

| | N | Mean | Median | SEM | z | p |
|---|---|---|---|---|---|---|
| HbR peak greatest locomotion | 21 | 0.947 | 0.948 | .006 | - | - |
| HbR peak least locomotion | 21 | 0.947 | 0.945 | .005 | - | - |
| Difference most vs least HbR | 21 | 0.000 | -.0004 | .005 | -.330 | .741 |

**Supplementary table S5: Sign test result for HbT and HbO greatest vs least locomotion (where locomotion is ranked across the entire 25s trial)**

| | N | Mean | Median | SEM | z | p |
|---|---|---|---|---|---|---|
| HbT peak greatest locomotion | 21 | 1.043 | 1.038 | .007 | - | - |
| HbT peak least locomotion | 21 | 1.030 | 1.028 | .003 | - | - |
| Difference most vs least HbT | 21 | .013 | .003 | .007 | -.873 | .383 |
| HbO peak greatest locomotion | 21 | 1.065 | 1.057 | .009 | - | - |
| HbO peak least locomotion | 21 | 1.053 | 1.049 | .005 | - | - |
| Difference most vs least HbO | 21 | .012 | -.0004 | .008 | .000 | 1.000 |

<u>Return to baseline</u>

**Supplementary table S6: Shapiro Wilk test of normality for baseline statistics (mean values taken from last 5s of the 2s whisker stimulation trial (between 15-20s)**

| Haemodynamic Measure | Statistic | df | p |
|---|---|---|---|
| HbT | .877 | 21 | .013 |
| HbO | .932 | 21 | .152 |
| HbR | .885 | 21 | .018 |

**Supplementary table S7: Sign test result for HbT and HbR return to baseline (HbT/HbR mean values between 15-20s for trials with the most locomotion compared with the mean values between 15-20s for trials with the least locomotion)**

| | N | Mean | Median | SEM | z | p |
|---|---|---|---|---|---|---|
| HbT most locomotion | 21 | 1.018 | 1.019 | .005 | - | - |
| HbT least locomotion | 21 | 0.995 | 0.997 | .002 | - | - |
| Difference HbT most vs least | 21 | .024 | .024 | .005 | -3.055 | .001 |
| HbR most locomotion | 21 | 0.982 | 0.980 | .004 | - | - |
| HbR least locomotion | 21 | 1.000 | 0.999 | .003 | - | - |
| Difference HbR most vs least | 21 | -.022 | -.021 | .005 | 3.491 | .000 |

**Supplementary table S8: Wilcoxon Signed Ranks test for HbO return to baseline (HbO mean value between 15-20s for trials with the most locomotion compared with the mean value between 15-20s for trials with the least locomotion)**

| | N | Mean | Median | SEM | z | p |
|---|---|---|---|---|---|---|
| HbO most locomotion | 21 | 1.028 | 1.030 | .006 | - | - |
| HbO least locomotion | 21 | 0.992 | 0.994 | .003 | - | - |
| Difference HbO most vs least | 21 | .036 | .032 | .007 | -3.702 | .000 |

Two-way Repeated measures ANOVAs

**Supplementary table S9: Shapiro Wilk test of normality for each time window, amount of locomotion and haemodynamic measures**

| | Most Locomotion | | | | | | Least Locomotion | | | | | |
|---|---|---|---|---|---|---|---|---|---|---|---|---|
| | **HbT** | | **HbO** | | **HbR** | | **HbT** | | **HbO** | | **HbR** | |
| **Time window** | df | p | df | p | df | p | df | p | df | p | df | p |
| -5-0 | 21 | .000* | 21 | .001* | 21 | .518 | 21 | .380 | 21 | .174 | 21 | .867 |
| 0-5 | 21 | .000* | 21 | .001* | 21 | .654 | 21 | .872 | 21 | .680 | 21 | .186 |
| 5-10 | 21 | .000* | 21 | .000* | 21 | .429 | 21 | .172 | 21 | .234 | 21 | .649 |
| 10-15 | 21 | .000* | 21 | .001* | 21 | .316 | 21 | .123 | 21 | .496 | 21 | .991 |
| 15-20 | 21 | .000* | 21 | .005* | 21 | .843 | 21 | .926 | 21 | .739 | 21 | .046* |

*Performed on studentised residuals. Time window is the time window in which locomotion was calculated for ranking purposes. *Indicates normality is violated*

**Supplementary table S10: Sphericity Results for the two-way repeated measures ANOVAs (Mauchly's test)**

| | **HbT** | | | | **HbO** | | | | **HbR** | | | |
|---|---|---|---|---|---|---|---|---|---|---|---|---|
| **Factor** | $X^2$ | df | p | ε | $X^2$ | df | p | ε | $X^2$ | df | p | ε |
| Time | 20.297 | 9 | .017 | .731 | 15.859 | 9 | .071 | .753 | 5.798 | 9 | .761 | .883 |
| Time * Locomotion | 24.286 | 9 | .004 | .645 | 17.771 | 9 | .039 | .723 | 20.637 | 9 | .015 | .632 |

*ε indicates Greenhouse Geiser value. $X^2$ Chi–squared value. Values <.05 sphericity violated*

**Supplementary table S11: Two-Way Repeated ANOVA outputs for HbT, HbO and HbR investigating the interaction between the amount of locomotion (factors: most & least) and the time at which locomotion occurred (factors: -5-0s, 0-5s, 5-10s, 10-15s, 15-20s)**

| | **HbT** | | | **HbO** | | | **HbR** | | |
|---|---|---|---|---|---|---|---|---|---|
| **Factor** | df | F | p | df | F | p | df | F | p |
| Time | 2.925, 58.492 | 7.849 | .000 | 4, 80 | 8.435 | .000 | 4, 80 | 9.134 | .000 |
| Locomotion | 1, 20 | 6.032 | .023 | 1, 20 | 4.868 | .039 | 1, 20 | 3.723 | .068 |
| Time * Locomotion | 2.578, 51.566 | 13.351 | .000 | 2.894, 57.876 | 13.317 | .000 | 2.526, 50.521 | 8.712 | .000 |

**Supplementary table S12: Sphericity Results (Mauchly's test) for simple main effects (One-way ANOVAs) investigating the effects of timing of locomotion on sensory-evoked hemodynamic responses for the most and least locomotion**

| | HbT | | | | HbO | | | | HbR | | | |
|---|---|---|---|---|---|---|---|---|---|---|---|---|
| | $X^2$ | df | p | ε | $X^2$ | df | p | ε | $X^2$ | df | p | ε |
| **Most** | | | | | | | | | | | | |
| Time | 29.923 | 9 | .000 | .630 | 30.653 | 9 | .000 | .665 | 19.251 | 9 | .024 | .672 |
| **Least** | | | | | | | | | | | | |
| Time | 17.153 | 9 | .047 | .735 | 11.929 | 9 | .219 | .772 | 13.192 | 9 | .156 | .729 |

*ε indicates Greenhouse Geiser value. Most refers to most locomotion, least refers to least locomotion*

**Supplementary table S13: Simple main effects to investigate the interaction between the amount and timing of locomotion and the effect on the haemodynamic response**

| **Factor: Time** | HbT | | | HbO | | | HbR | | |
|---|---|---|---|---|---|---|---|---|---|
| | df | F | p | df | F | p | df | F | p |
| **Most Locomotion** | 2.520, 50.409 | 12.998 | .000 | 2.660, 53.192 | 12.788 | .000 | 2.689, 53.781 | 11.504 | .000 |
| **Least Locomotion** | 2.940, 58.791 | 8.933 | .000 | 4, 80 | 9.939 | .000 | 4, 80 | 6.829 | .000 |

*One-way ANOVAs were used to assess simple main effects

**Supplementary table S14: Notable comparisons between 5s time windows for HbT, HbO and HbR peaks during whisker stimulation during most or least locomotion conditions**

| Haemodynamic Measure | Time window for ranking locomotion and time window compared to | P value |
|---|---|---|
| **Most Locomotion** | | |
| HbT | -5-0 > 15-20 | .013 |
| | 0-5 > 5-10 | .006 |
| | 0-5 > 10-15 | .003 |
| | 0-5 > 15-20 | .000 |
| HbO | -5-0 > 10-15 | .048 |
| | -5-0 > 15-20 | .008 |
| | 0-5 > 5-10 | .008 |
| | 0-5 > 10-15 | .003 |
| | 0-5 > 15-20 | .000 |
| HbR | -5-0 < 5-10 | .017 |
| | -5-0 < 10-15 | .012 |
| | -5-0 < 15-20 | .005 |
| | 0-5 < 5-10 | .013 |
| | 0-5 < 10-15 | .004 |
| | 0-5 < 15-20 | .001 |
| **Least Locomotion** | | |
| HbT | -5-0 > 0-5 | .000 |
| | 0-5 < 5-10 | .013 |
| | 0-5 < 10-15 | .009 |
| | 0-5 < 15-20 | .005 |
| HbO | -5-0 > 0-5 | .000 |
| | 0-5 < 5-10 | .006 |
| | 0-5 < 10-15 | .001 |
| | 0-5 < 15-20 | .002 |
| HbR | -5-0 < 0-5 | .008 |
| | 0-5 > 5-10 | .018 |
| | 0-5 > 10-15 | .007 |
| | 0-5 > 15-20 | .015 |

*> Indicates greater than and < indicates less than*

**Supplementary table S15: Pairwise comparisons with a Bonferroni correction comparing mean peaks for trials in which most and least locomotion occurred during the different time windows**

| | | **Most Locomotion** | | **Least Locomotion** | | | | |
|---|---|---|---|---|---|---|---|---|
| **Time window** | Measure | *M* | *SEM* | *M* | *SEM* | *df* | *F* | *p* |
| -5-0 | HbT | 1.049 | ± .008 | 1.035 | ± .003 | 1, 20 | 3.466 | .077 |
| | HbO | 1.078 | ± .011 | 1.061 | ± .006 | 1, 20 | 2.401 | .137 |
| | HbR | 0.930 | ± .008 | 0.944 | ± .006 | 1, 20 | 2.149 | .158 |
| 0-5 | HbT | 1.054 | ± .007 | 1.020 | ± .002 | 1, 20 | 19.676 | .000* |
| | HbO | 1.083 | ± .009 | 1.033 | ± .004 | 1, 20 | 24.825 | .000* |
| | HbR | 0.934 | ± .006 | 0.969 | ± .004 | 1, 20 | 38.753 | .000* |
| 5-10 | HbT | 1.041 | ± .007 | 1.031 | ± .003 | 1, 20 | 3.144 | .091 |
| | HbO | 1.063 | ± .009 | 1.052 | ± .005 | 1, 20 | 1.702 | .207 |
| | HbR | 0.948 | ± .005 | 0.952 | ± .005 | 1, 20 | .486 | .494 |
| 10-15 | HbT | 1.037 | ± .005 | 1.034 | ± .004 | 1, 20 | .756 | .395 |
| | HbO | 1.058 | ± .007 | 1.057 | ± .006 | 1, 20 | .057 | .814 |
| | HbR | 0.951 | ± .005 | 0.948 | ± .006 | 1, 20 | .316 | .580 |
| 15-20 | HbT | 1.032 | ± .004 | 1.031 | ± .003 | 1, 20 | .058 | .812 |
| | HbO | 1.052 | ± .006 | 1.052 | ± .005 | 1, 20 | .000 | .995 |
| | HbR | 0.954 | ± .005 | 0.951 | ± .005 | 1, 20 | .219 | .645 |

**Indicates a significant difference between the mean peak of the haemodynamic measure when comparing most and least locomotion, with Bonferroni adjustment applied. M indicates mean and SEM indicates standard error of the mean*
